# OMGS: Optical Map-based Genome Scaffolding

**DOI:** 10.1101/585794

**Authors:** Weihua Pan, Tao Jiang, Stefano Lonardi

## Abstract

Due to the current limitations of sequencing technologies, *de novo* genome assembly is typically carried out in two stages, namely contig (sequence) assembly and scaffolding. While scaffolding is computationally easier than sequence assembly, the scaffolding problem can be challenging due to the high repetitive content of eukaryotic genomes, possible mis-joins in assembled contigs and inaccuracies in the linkage information. Genome scaffolding tools either use paired-end/mate-pair/linked/Hi-C reads or genome-wide maps (optical, physical or genetic) as linkage information. Optical maps (in particular Bionano Genomics maps) have been extensively used in many recent large-scale genome assembly projects (e.g., goat, apple, barley, maize, quinoa, sea bass, among others). However, the most commonly used scaffolding tools have a serious limitation: they can only deal with one optical map at a time, forcing users to alternate or iterate over multiple maps. In this paper, we introduce a novel scaffolding algorithm called OMGS that for the first time can take advantages of multiple optical maps. OMGS solves several optimization problems to generate scaffolds with optimal contiguity and correctness. Extensive experimental results demonstrate that our tool outperforms existing methods when multiple optical maps are available, and produces comparable scaffolds using a single optical map. OMGS can be obtained from https://github.com/ucrbioinfo/OMGS

## 1 Introduction

Genome assembly is a fundamental problem in genomics and computational biology. Due to the current limitations of sequencing technologies, the assembly is typically carried out in two stages, namely contig (sequence) assembly and scaffolding. Scaffolds are arrangements of oriented contigs with gaps representing the estimated distance separating them. The scaffolding process can vastly improve the assembly contiguity and can produce chromosome-level assemblies. Despite significant algorithmic progress, the scaffolding problem can be challenging due to the high repetitive content of eukaryotic genomes, possible mis-joins in assembled contigs and the inaccuracies of the linkage information.

Genome scaffolding tools either use paired-end/mate-pair/linked/Hi-C reads or genome-wide maps. The first group includes scaffolding tools for second generation sequencing data, such as Bambus [29, 17], GRASS [13], MIP [31], Opera [12], SCARPA [11], SOPRA [8] and SSPACE [5] and the scaffolding modules from assemblers ABySS [35], SGA [34] and SOAPdenovo2 [22]. Since the relative orientation and approximate distance between paired-end/mate-pair/linked/Hi-C reads are known, the consistent alignment of a sufficient number of reads to two contigs can indicate their relative order, their orientation and the distance between them. An extensive comparison of scaffolding methods in this first group of tools can be found in [14].

The second group uses genome-wide maps such as genetic maps [37], physical maps, or optical maps. According to the markers provided by these maps, contigs can be anchored to specific positions so that their order and orientations can be determined. The distance between contigs can also be estimated with varying degree of accuracy depending on the density of the map.

The optical mapping technologies currently on the market (e.g., BioNano Genomics Irys systems, OpGen Argus) allow computational biologists to produce genome-wide maps by fingerprinting long DNA molecules (up to 1 Mb), via nicking restriction enzymes [32]. Linear DNA fragments are stretched on a glass surface or in a nano-channel array, then the locations of restriction sites are identified with the help of dyes or fluorescent labels. The results are imaged and aligned to each other to map the locations of the restriction sites relative to each other. While the assembly process for optical molecules is highly reliable, there is clear evidence that a small fraction of the optical molecules is chimeric [15].

A few scaffolding algorithms that use optical maps are available. SOMA appears to be the first published tool that can take advantage of optical maps but it can only deal with a non-fragmented optical map [25]. The scaffolding tool proposed in [30] was used for two bacterial genomes *Yersinia pestis* and *Yersinia enterocolitica*, but the software is no longer publicly available. In the last few years, Bionano optical maps have become very popular, and have been used to improve the assembly contiguity in many large-scale *de novo* genome assembly projects (e.g., goat, apple, barley, maize, quinoa, sea bass [4, 28, 7, 23]). To the best of our knowledge, the main tools used to generate scaffolds using Bionano optical maps are SewingMachine from KSU [33] and HybridScaffold from Bionano Genomics (unpublished, 2016). SewingMachine seems to be favored by practitioners over HybridScaffold.

Both HybridScaffold and SewingMachine have, however, a serious limitation: they can only deal with one optical map at a time, forcing users to alternate or iterate over optical maps when multiple maps are available. In this paper, we introduce a novel scaffolding algorithm called OMGS that for the first time can take advantage of any number of optical maps. OMGS solves several optimization problems to generate scaffolds with optimal contiguity and correctness.

## 2. Problem definition

The input to the problem is the genome assembly to be scaffolded (represented by a set of assembled contigs), and one or more optical maps (represented by a set of sets of genomic distances). We use *C* = {*c*_*i*_|*i* = 1, …, *l*} to denote the set of contigs in the genome assembly, where each *c*_*i*_ is a string over the alphabet {*A, C, G, T*}. Henceforth, we assume that the contigs in *C* are chimera-free.

An optical map is composed by a set of optical molecules, each of which is represented by an ordered set of positions for the restriction enzyme sites. As said, optical molecules are obtained by an assembly process similar to sequence assembly, but we will reserve the term *contig* for sequenced contigs. We use *M* = {*m*_*i*_|*i* = 1, …, *n*} to denote the optical map, where each optical molecule *m*_*i*_ is an ordered set of integers, corresponding to the distances in base pairs between two adjacent restriction enzyme sites on molecule *m*_*i*_. By digesting *in silico* the contigs in *C* using the same restriction enzyme used to produce the optical map and matching the sequence of adjacent distances between sites, one can align the contigs in *C* to the optical map *M*. If one is given multiple optical maps obtained using different restriction enzymes, *M* will be the union of the molecules from all optical maps. In this case, each genomic location is expected to be covered by multiple molecules in *M*. As said, high quality alignments allows one to anchor and orient contigs to specific coordinates on the optical map. When multiple contigs align to the same optical map molecule, one can order them and estimate the distance between them. By filling these gaps with a number of *N*’s equal to the estimated distance, longer DNA sequences called *scaffolds* can be obtained.

A series of practical factors make the problem of scaffolding non-trivial. These factors include imprecisions in optical maps (e.g., mis-joins introduced during the assembly of the optical map [15]), unreliable alignments between contigs and optical molecules, and multiple inconsistent anchoring positions for the same contigs. As a consequence, it is appropriate to frame this scaffolding problem as an optimization problem.

We are now ready to define the problem. We are given an assembly represented by a set of contigs *C*, a set of optical map molecules *M* and a set of alignments *A* = {*a*_1,1_, *a*_1,2_, *… a*_*l,n*_} of *C* to *M*, where *a*_*i,j*_ is the alignment of contig *c*_*i*_ to optical map molecule *o*_*j*_. The problem is to obtain a set of scaffolds *S* = {*s*_1_, *s*_2_, *… s*_*k*_} where each *s*_*i*_ is a string over the alphabet {*A, C, G, T, N*}, such that (i) each contig *c*_*i*_ is contained/assigned to exactly one scaffold, (ii) the *contiguity* of *S* is maximized and (iii) the conflicts of *S* with respect to *A* are minimized. This optimization problem is not rigorously defined unless one defines precisely the concepts of *contiguity* and *conflict*, but this description captures the spirit of what we want to accomplish. In genome assembly, the assembly contiguity is usually captured by statistical measures like the N50/L50 or the NG50/LG50. The notion of conflict is not easily quantified, and even if it was made precise, this multi-objective optimization problem would be hard to solve. We decompose this problem into two separate steps, namely (a) scaffold detection and (b) gap estimation, as explained below.

## 3 Method

As said, our proposed method is composed of two phases: scaffold detection and gap estimation. In the first phase, contigs are grouped into scaffolds and the order of contigs in each scaffold is determined. In the second phase, distances between neighboring contigs assigned to scaffolds are estimated. The pipeline of the proposed algorithm is illustrated in Figure 1.

**Figure 1:**
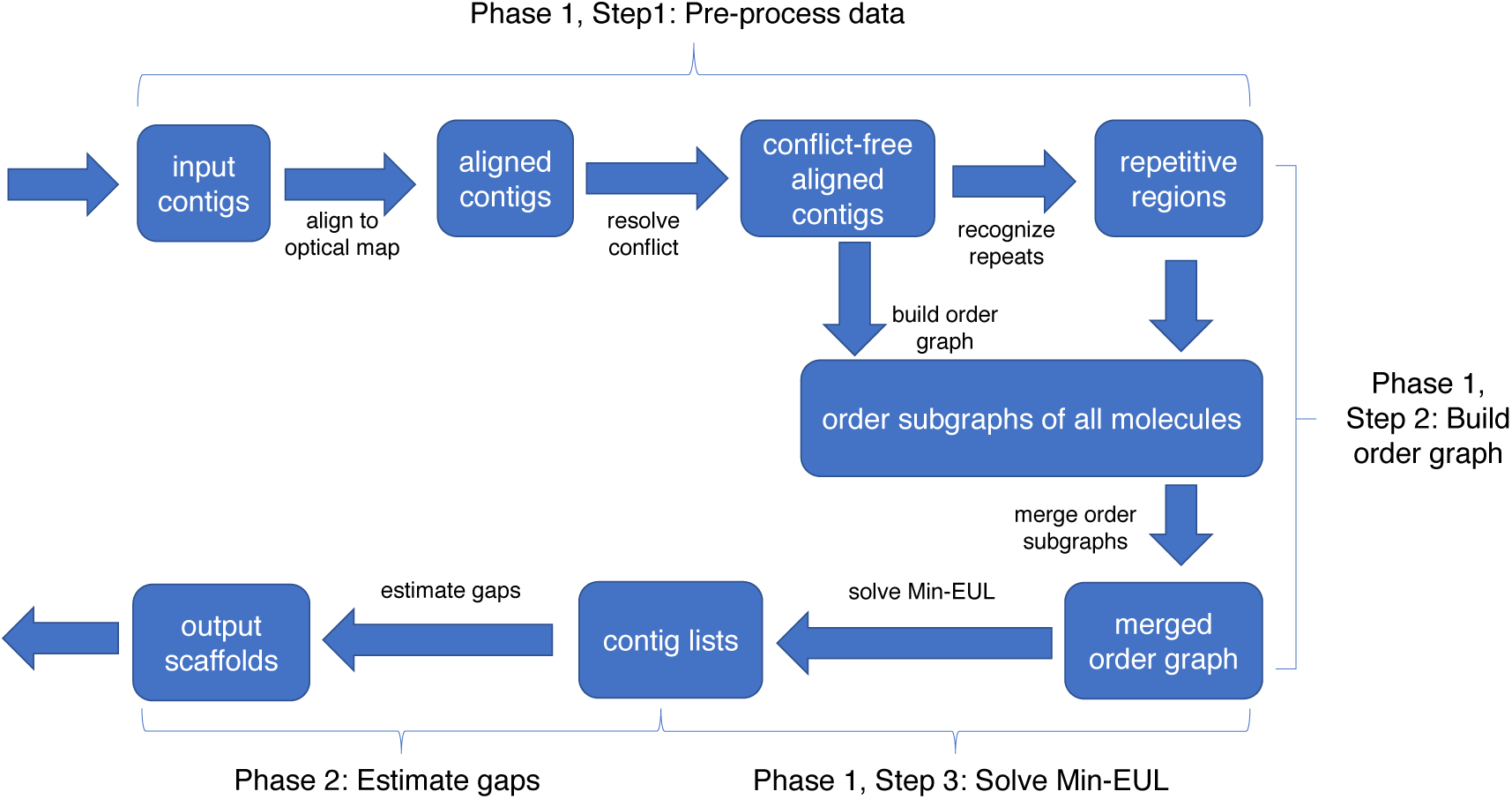
Pipeline of the proposed algorithm

### 3.1 Phase 1: Detecting scaffolds

Phase 1 has three major steps. In Step 1, we align *in silico*-digested chimeric-free contigs to the optical maps (e.g., for a Bionano optical map, we use RefAligner), but not all alignments are used in Step 2. We only consider alignments that (i) exceed a minimum confidence level (e.g, confidence 15 in the case of RefAligner); (ii) do not overlap each other more than a given genomic distance (e.g, 20 kbp) and (iii) do not create conflict with each other. The method we use here to select conflict-free alignments was introduced in our previous work [27]. In Step 2, we compute candidate scaffolds by building the *order graph* and formulating an optimization problem on it. In Step 3, either the exhaustive algorithm or a log *n*-approximation algorithm is used to solve the optimization problem (depending on the size of the graph) and produce the final scaffolds.

#### 3.1.1 Building the order graph

The order graph *O* is a directed weighted graph in which each vertex represents a contig. Given two contigs *c*_*i*_ and *c*_*j*_ aligned to an optical molecule *o* with alignments *a*_*i*_ and *a*_*j*_, we create a directed edge (*c*_*i*_, *c*_*j*_) in *O* if (i) the starting coordinate of alignment *a*_*i*_ (that we call *a*_*i*_.*start* henceforth) is smaller than the starting coordinate of alignment *a*_*j*_ (that we call *a*_*j*_. *start* henceforth) and (ii) there is no other alignment *a*_*k*_ such that *a*_*k*_.*start* is between *a*_*i*_.*start* and *a*_*j*_.*start* and (iii) there are no conflict sites between *a*_*i*_.*end* and *a*_*j*_.*start* on the optical molecule, as defined below. For each alignment *a* between optical molecule *o* and contig *c*, we compute the left overhang *l*_*o*_ and right overhang *r*_*o*_ from *o* and the left overhang *l*_*c*_ and right overhang *r*_*c*_ from *c*. The left-end of alignment *a* is declared a *conflict site* if (i) both *l*_*o*_ and *l*_*c*_ are longer than some minimum length (e.g., 50 kbp) and (ii) at least one restriction enzyme sites appear in both *l*_*o*_ and *l*_*c*_. A symmetric argument applies to the right-end of the alignment, which determines the values for *r*_*o*_ and *r*_*c*_.

Directed edge (*c*_*i*_, *c*_*j*_) is assigned a weight equal to qual(*o, a*_*i*_.*end, a*_*j*_.*start*) * (conf(*a*_*i*_)+ conf(*a*_*j*_)), where (i) qual(*o, a*_*i*_.*end, a*_*j*_.*start*) is the *quality* of the region between *a*_*i*_.*end* and *a*_*j*_.*start* on molecule *o* (higher is better, defined next) and (ii) conf(*a*) is the confidence score provided by RefAligner alignment *a* (higher is better). The quantity qual(*o, s, t*) is defined based on the length of a repetitive region between coordinates (*s, t*). Based on our experience, assembly mis-joins on optical molecule almost always happen in repetitive regions [15]. Given the length of repetitive region len rep(*o, s, t*) in base pairs (defined below), we define the quality of *o* in the interval (*s, t*) as qual(*o, s, t*) = *e*^−len_rep(*o,s,t*)*/*100000^. When *a*_*i*_ and *a*_*j*_ have a small overlap (e.g., shorter than 20 kbp), we set len rep(*o, s, t*) = 0.

We recognize repetitive regions in optical molecules based on the distribution of restriction enzyme sites. For a molecule *o* with *n* sites, let *m*_*i*_ be the coordinate of the *i*-th site for *i* = 1, *…, n*. As said, molecule *o* can be represented as a list of positions {*m*_*i*_|*i* = 1, *…, n*}. In order to determine the repetitive regions in *o*, we slide a window that covers *k* sites (e.g., *k* = 10 sites). At each position *j* = 1, *…, n − k* + 1, we select window *w*_*j*_ = {*m*_*j*_, *…, m*_*j*+*k-*1_}. While repetitive regions in genome can be highly complex (see, e.g., [40]), we observed only two types of repetitive regions in optical molecules, namely single-site repetitive region (see Figure 2-A) and two-site repetitive region (see Figure 2-B). It is entirely possible that more complex repetitive regions exist: if they do, they seem rare. Based on this observation, in order to decide whether window *w*_*j*_ is repetitive, we first compute two lists of pairwise distances between sites, namely *D*_*j*,1_ = {*m*_*j*+*l*_ − *m*_*j*+*l*−1_|*l* = 1, …, *k* − 1} and *D*_*j*,2_ = {*m*_*j*+*l*+1_ − *m*_*j*+*l* − 1_|*l* = 1, …, *k* − 2} that we call *distance lists*, then we apply the statistical test described next.

**Figure 2:**
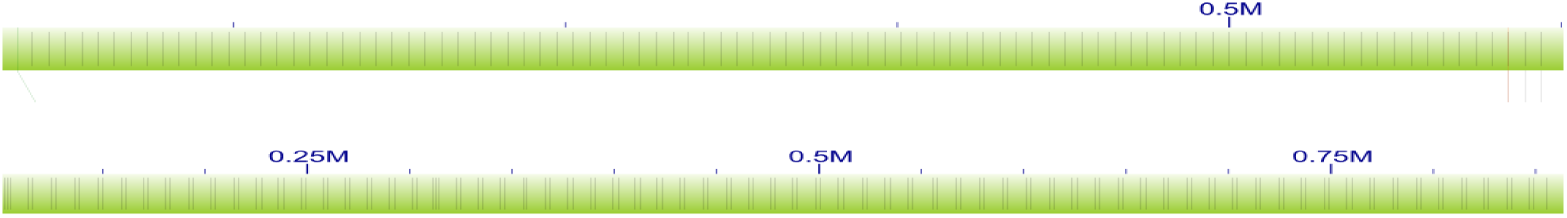
Examples of single-site repetitive region (A) and two-site repetitive region (B) in optical maps. Observe the small variations in the repetitive patterns in (B)

In our statistical test we assume that the values in the distance lists that belong to repetitive regions are independent and identically distributed as a Gaussian. We further assume that each specific distance list (*D*_*j*,1_ or *D*_*j*,2_) is associated with a Gaussian with a specific mean *µ*_*j,q*_ (*q ∈ {*1, 2}). Finally, we assume that the variance *σ* is globally shared by all molecules. An estimator of the mean is *µ*_*j,q*_ is 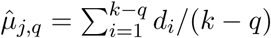, where *d*_*i*_ *∈ D*_*j,q*_ and *k* is the window size. To estimate *σ*^2^, we first get an initial (rough) estimate of the repetitive regions on all molecules. Given a particular *D*_*j,q*_, let *d*_*max*_ and *d*_*min*_ be the maximum and minimum distance in *D*_*j,q*_. We declare a distance list *D*_*j,q*_ to be *estimated repetitive* if *d*_*max*_ − *d*_*min*_ is smaller than a given distance (e.g., 1.5 kbp). We collect all estimated repetitive lists in set *R* = {*D*_*p*_ is estimated repetitive|*p* = 1, *…, P*} and the estimated mean 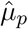 for each distance list *D*_*p*_ in the set *R*, where *P* is the total number of estimated repetitive lists. Then, we define the log likelihood function *L* as follows (additional details can be found in Appendix, Section B)

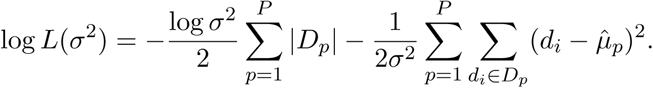

By maximizing log *L*(*σ*^2^), the estimator for the variance becomes

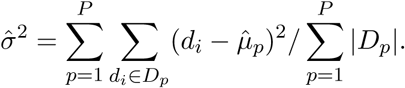

Then, we carry out the test on the statistic *d*_*max*_ − *d*_*min*_ for each *D*_*j,q*_. The joint density function of (*d*_*max*_, *d*_*min*_) is

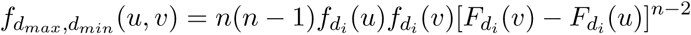

for *-∞ < u < v <* +*∞*, where 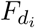 and 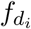 are the distribution function and density function of 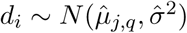, respectively. The density function of *d*_*max*_ − *d*_*min*_ is

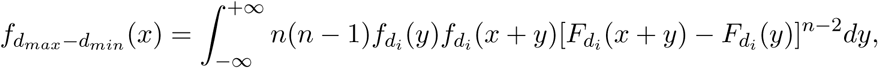

defined when *x* ≥ 0 (additional details can be found in Appendix, Section C). Let now *X* be a random variable associated with the distribution 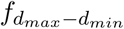. If the p-value *P* (*X > d*_*max*_ − *d*_*min*_) is greater than a predefined threshold (e.g., 0.001), we accept the null hypothesis and declare that window *w*_*j*_ is repetitive. The repetitive regions for the entire molecule *o* is the union of all the windows *w*_*j*_’s recognized as repetitive according to the test above.

Once the order graph of each optical molecule is built, we connect all the order graphs which share the same contigs using the association graph introduced in [27]. The association graph is an undirected graph in which each vertex represents an optical molecule and an edge indicates that the two molecules share at least one contig aligned to both of them. We use depth first search (DFS) to first build a spanning forest of the association graph. Then, we traverse each spanning tree and connect the corresponding order subgraph to the final order graph. Every time we add a new graph, new vertices and new edges might be added. If an edge already exist, the weights of the new edges are added to the weights of existing edges.

#### 3.1.2 Generating scaffolds

Once the order graph *O* is finalized, we generate the ordered sequence of contigs in each scaffold. In the ideal case, each connected component *O*_*i*_ of *O* is a directed acyclic graph (DAG) because the genome is one-dimensional and the order of any pair of contigs is unique. In practice however, *O*_*i*_ may contain cycles caused by the inaccuracy of the alignments and mis-joins in optical molecules. To convert each cyclic component *O*_*i*_ into a DAG, we solve the MINIMUM FEEDBACK ARC SET problem on *O*_*i*_. In this problem, the objective is to find the minimum subset of edges (called *feedback arc set*) containing at least one edge of every cycle in the input graph. Since the minimum feedback edge set problem is APX-hard, we use the greedy local heuristics introduced in [2] to solve it.

We then break each DAG *G*_*i*_ of connected component *O*_*i*_ into subgraphs as follows. In each subgraph, we require that the order of every pair of vertices to be uniquely determined by the directed edges. This allows us to uniquely determine the order of the contigs for each scaffold. The formal definition of this optimization problem is as follows.

##### Definition 1

(Minimum Edge Unique Linearization problem). Input: A weighted directed acyclic graph *G* = (*V, E*). Output: A subset of edges *E′ ⊆ E* such that (i) in each connected component 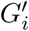 of the graph *G′* = (*V, E* − *E′*) obtained after removing *E′*, the order of all vertices can be uniquely determined, and (ii) the total weights of the edges in *E′* is the minimum among all the subset of edges satisfying (i).

In Theorem 1 below, we show that the Minimum Edge Unique Linearization problem (Min-EUL) is NP-hard by proving that it is equivalent to the Minimum Edge Clique Partition problem (Min-ECP), which is know to be NP-hard [10]. In Min-ECP, we are given a general undirected graph, and we need to partition its vertices into disjoint clusters such that each cluster forms a clique and the total weight of the edges between clusters is minimized.

##### Theorem 1.

Min-EUL *is equivalent to* Min-ECP.

*Proof.* First, we show that Min-EUL polynomially reduces to Min-ECP. Given an instance *G* = (*V, E*) of Min-EUL, we build an instance *G′* = (*V ′, E′*) of Min-ECP as follows. Let *V ′* = *V*. For each pair of vertices *u, v ∈ V ′* where *v* is reachable from *u*, define an undirected edge between *u* and *v* in *E′*. For each directed edge (*u, v*) *∈ E*, set the weight of the corresponding undirected edge (*u, v*) *E′* as 1. Set the weights of the other edges in *E′* as 0. Then it is easy to see that a Min-EUL solution to *G′* is equivalent to a Min-ECP solution to *G* and vice versa.

Now we show that Min-ECP polynomially reduces to Min-EUL. Given an instance *G′* = (*V ′, E′*) (assuming *G′* is connected) of Min-ECP, we build an instance *G* = (*V, E*) of Min-EUL as follows. Let *V* = *V ′*. Pick any total linear order *O* of all vertices in *V ′*. For each undirected edge (*u, v*) *∈ E′* where rank(*u*) *<* rank(*v*) in *O*, define a directed edge from *u* to *v* in *E* and set its weight to be the same as its corresponding undirected edge in *E′*. For any two vertices *u, v ∈ V*, where rank(*u*) *<* rank(*v*) and (*u, v*) *∉ E′*, add a new vertex *x*_*uv*_ *∈ V* with rank(*x*_*uv*_) = rank(*v*) and a directed edge *u* to *x*_*uv*_ of weight 1 in *E*. Now for each pair of vertices *u, v ∈ V* where rank(*u*) *<* rank(*v*) and (*u, v*) *∉ E*, add a directed edge *u* to *v* with weight zero in *E*. Then it is easy to see that a Min-EUL solution to *G* corresponds to a Min-ECP solution to *G* and vice versa.

Given the complexity of Min-EUL, we propose an exponential time exact algorithm and a polynomial time log *n*-approximation algorithm for solving it. To describe the exact algorithm, we need to introduce some notations. A *conjunction* vertex in a DAG is a vertex which has more than one incoming edge or outgoing edge. A *candidate* edge is an edge which connects at least one conjunction vertex. In Theorem 2 below, we prove that the optimal solution *E′* of Min-EUL must only contain candidate edges. Let *E*_*c*_ be the set of all candidate edges in the DAG *G*, for each subset 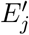 of *E*_*c*_, we check whether the graph 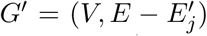 satisfies requirement (i) in Definition 1 after removing 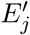 from *G*. Among all the feasible 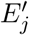, we produce the set of edges with minimum total weights. To check whether 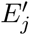 is feasible, we use a variant of topological sorting which requires one to produce a unique topological ordering. To do so, we require that in every iteration of topological sorting, the candidate node to be added to sorted graph is always unique. Details of this algorithm are shown as Algorithm 1 in Appendix A.

##### Theorem 2.

*The optimal solution E′ of* Min-EUL *only contains candidate edges.*

*Proof.* For sake of contradiction, we assume that *E′* contains a non-candidate edges (*u, v*). Since *E′* is optimal, *G′* = (*V, E − E′*) satisfies condition (i) in Definition 1. Since both *u* and *v* are conjunction vertices, *u* has only one incoming edge and *v* has only one outgoing edge. Therefore, by adding (*u, v*) to *G′* = (*V, E − E′*), we still satisfy condition (i) in Definition 1. Since the weight of (*u, v*) is positive, the total weight of *E − E′* + {(*u, v*)} is larger than *E − E′*. Therefore *E′ − {*(*u, v*)} is optimal, contradicting the optimality of *E′*.

As said, Min-EUL is equivalent to Min-ECP (Theorem 1). In addition, the authors of [10] showed that for any instance of Min-ECP one can find an equivalent instance of the Minimum Disagreement Correlation Clustering problem. As a consequence, any algorithm for the Minimum Disagreement Correlation Clustering problem could be used to solve Min-EUL. In our tool OMGS, we implemented a *O*(log *n*)-approximation algorithm based on linear programming, originally proposed in [9]. Standard linear programming packages (e.g., GLPK or CPLEX) are used to solve the linear program. We use the exact algorithm for DAGs with no more than twenty candidate edges, and the approximation algorithm for larger DAGs.

### 3.2 Phase 2: Estimating gaps

Let *s* = {*c*_*i*_|*i* = 1, *…, h*} be one of the scaffold generated in Phase 1 where each *c*_*i*_ is a contig. In Phase 2, we estimate the length *l*_*i*_ of the gap between each pair *c*_*i*_ and *c*_*i*+1_ of adjacent contigs. We estimate all gap lengths *L* = {*l*_*i*_|*i* = 1, *…, h -* 1} at the same time using the distances between the contigs provided by the alignments and the corresponding order subgraphs. We assume that each *l*_*i*_ is chi-square distributed with *α*_*i*_ degrees of freedom. The choice of chi-square distribution is due to its additive properties, namely the sum of independent chi-squared variables is also chi-squared distributed. Recall that each order subgraph *O*_*k*_ provides an unique ordering *x*_*k*_ = {*c*_*j*_|*j* = 1, *…, r*} of the contigs aligned to molecule *o*_*k*_, while the coordinates of the alignment provide the distances between all pairs of adjacent contigs *c*_*j*_ and *c*_*j*+1_ as *y*_*k*_ = {*d*_*j*_|*j* = 1, *…, r -* 1}. We use the distances *d*_*j*_ as samples to estimate gap lengths *l*_*i*_. If edge (*c*_*j*_, *c*_*j*+1_) in *O*_*k*_ is removed in the order graph *O* when solving Min-EUL in Phase 1, *d*_*j*_ will be considered not reliable and removed from *y*_*k*_.

In the ideal case, *d*_*j*_ should be a sample of a single *l*_*i*_ (i.e., *c*_*j*_*c*_*j*+1_ in *x*_*k*_ corresponds to *c*_*p*_*c*_*p*+1_ in *s*). In practice however, *c*_*j*_*c*_*j*+1_ in *x*_*k*_ will corresponds to a different pair *c*_*p*_*c*_*q*_ in *s* where *q > p* + 1 (i.e., *c*_*p*+1_ *… c*_*q-*1_ are missing from the order subgraph because some alignments with low confidence were removed in Step 1 of Phase 1). In this situation, after subtracting the length of missing contigs from 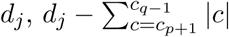 is a sample of 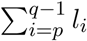 where |*c*| represents the length of contig *c*. Since *l*_*p*_, *…, l*_*q-*1_ are independent chi-square random variables, 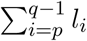 is chi-square distributed with degree of freedom 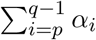, so that the log likelihood of this sample is

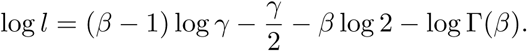

where 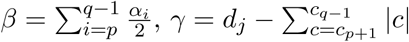 and Γ is the gamma function (additional details can be found in Appendix, Section D). The total log likelihood is the sum of the log likelihoods across all samples. To find the *α*_*i*_ maximizing the total log likelihood, we use the Broyden-Fletcher-Goldfarb-Shanno (BFGS) algorithm [1]. Since the mean of a chi-square distribution equals its degree of freedom, we obtain the estimated gaps 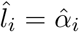. For the case in which the *l*_*i*_ are pre-estimated as negative in the first step, the second and third steps are ignored and the pre-estimated distances are used as final estimates.

Finally, we add 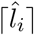 nucleotides (represented by Ns) between each pair of contigs *c*_*i*_ and *c*_*i*+1_. When 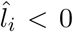, we add exactly 100 Ns between *c*_*i*_ and *c*_*i*+1_, which is the convention for a gap of unknown length.

## 4 Experimental results

We compared OMGS against KSU SewingMachine (version 1.0.6, released in 2015) and Bionano HybridScaffold (version 4741, released in 2016) which, to the best of our knowledge, are the only available scaffolding tools for Bionano Genomics optical maps. All tools were run with default parameters, unless otherwise specified. We collected experimental results on scaffolds of (i) cowpea (*Vigna unguiculata*) and (ii) fruit fly (*Drosophila melanogaster*).

### 4.1 Experimental results on cowpea

Cowpea is a diploid with a chromosome number 2*n* = 22 and an estimated genome size of 620 Mb. We sequenced the cowpea genome using single-molecule real-time sequencing (Pacific Biosciences RSII). A total of 87 SMRT cells yielded about 6M reads for a total of 56.84 Gbp (91.7x genome equivalent). We tested the three scaffolding tool on a high-quality assembly produced by Canu [3, 18] with parameters `corMhapSensitivity=high` and `corOutCoverage=100`, then polished it with Quiver. We used Chimericognizer to detect and break chimeric contigs, using seven other assemblies generated by Canu, Falcon [6] and ABruijn [20] as explained in [26].

In addition to standard contiguity statistics (N50^1^, L50^2^), total assembled size and scaffold length distribution, we determined incorrect/chimeric scaffolds by comparing them against the high-density genetic map available from [24]. We BLASTed 121bp-long design sequence for the 51,128 genome-wide SNPs described in [24] against each assembly, then we identified which contigs had SNPs mapped to them, and what linkage group (chromosome) of the genetic map those mapped SNPs belonged to. Chimeric contigs were revealed when their mapped SNPs belonged to more than one linkage group. The last line of Table 1 and Table 2 report the total size of contigs in each assembly for which (i) they have at least one SNPs mapped to it and (ii) all SNPs belong to the same linkage group (i.e., likely to be non-chimeric).

**Table 1:**
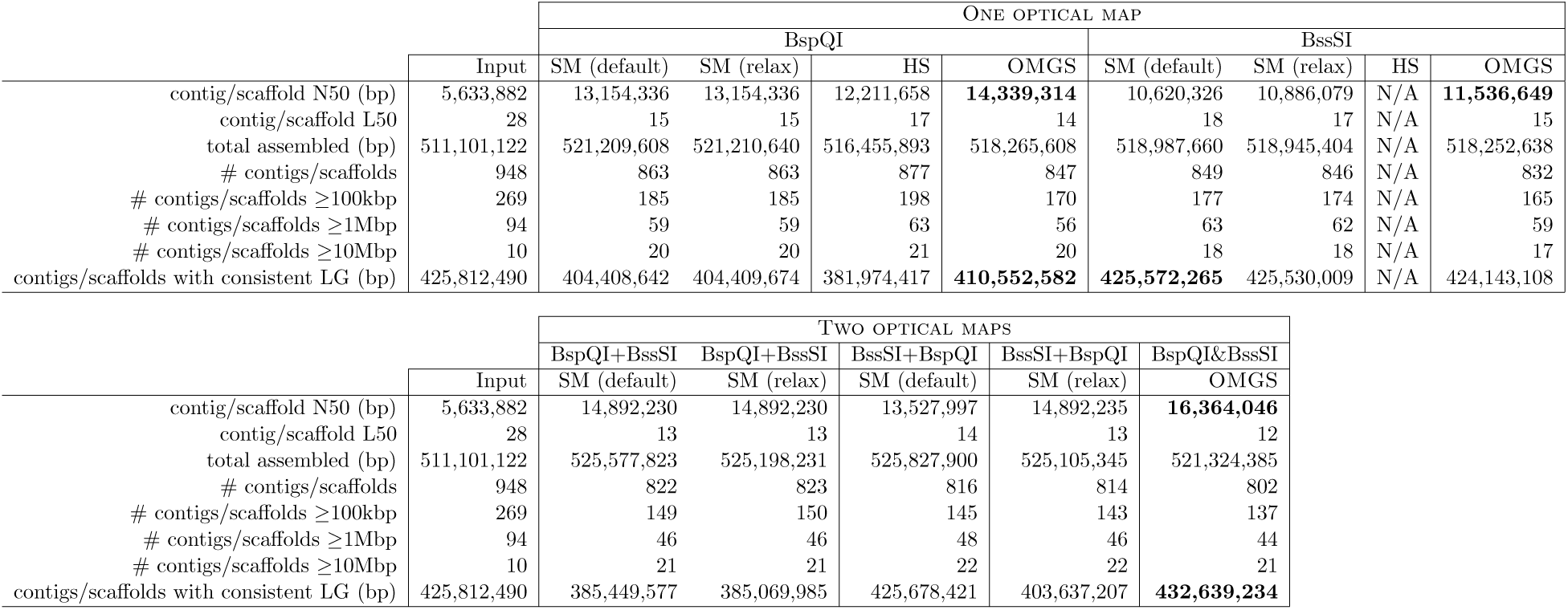
Comparing OMGS, SewingMachine (SM) and HybridScaffold (HS) on a cowpea assembly using one or two optical maps. Numbers in boldface highlight the best N50 and scaffold consistency with the genetic map for one map (BspQI and BssSI) or two maps (‘A+B’ refers to the use of map A followed by map B, ‘A&B’ refers to the use of both maps at the same time).

**Table 2:** Comparing OMGS, SewingMachine (SM) and HybridScaffold (HS) on a cowpea assembly using optical maps corrected by Chimericognizer. Numbers in boldface highlight the best N50 and scaffold consistency with the genetic map for one map (BspQI and BssSI) or two maps (‘A+B’ refers to the use of map A followed by map B, ‘A&B’ refers to the use of both maps at the same time).

|  |  | ONE OPTICAL MAP |  |  |  |  |  |  |  |
| --- | --- | --- | --- | --- | --- | --- | --- | --- | --- |
|  |  | BspQI |  |  |  | BssSI |  |  |  |
|  | Input | SM (default) | SM (relax) | HS | OMGS | SM (default) | SM (relax) | HS | OMGS |
| contig/scaffold N50 (bp) | 5,633,882 | 12,487,373 | 12,487,373 | 12,495,655 | **13,505,314** | 9,420,899 | 10,886,079 | N/A | **11,256,770** |
| contig/scaffold L50 | 28 | 16 | 16 | 15 | 14 | 19 | 17 | N/A | 16 |
| total assembled (bp) | 511,101,122 | 519,785,777 | 519,785,777 | 515,519,585 | 518,405,022 | 517,678,278 | 517,636,022 | N/A | 517,318,151 |
| # contigs/scaffolds | 948 | 863 | 863 | 871 | 849 | 854 | 851 | N/A | 837 |
| # contigs/scaffolds $\geq 100$kb | 269 | 185 | 185 | 192 | 172 | 182 | 179 | N/A | 169 |
| # contigs/scaffolds $\geq 1$Mbp | 94 | 60 | 60 | 60 | 58 | 66 | 65 | N/A | 62 |
| # contigs/scaffolds $\geq 10$Mbp | 10 | 19 | 19 | 19 | 19 | 17 | 17 | N/A | 17 |
| contigs/scaffolds with consistent LG (bp) | 425,812,490 | 413,819,557 | 413,819,557 | 402,840,302 | **421,466,164** | **424,262,883** | 424,220,627 | N/A | 423,117,331 |

**Table 2:** Comparing OMGS, SEWINGMACHINE (SM) and HYBRIDSCAFFOLD (HS) on a cowpea assembly using optical maps corrected by CHIMERICOGNIZER.
|  |  | TWO OPTICAL MAPS |  |  |  |  |  |
| --- | --- | --- | --- | --- | --- | --- | --- |
|  |  | BspQI+BssSI | BspQI+BssSI | BssSI+BspQI | BssSI+BspQI | BspQI&BssSI |  |
|  | Input | SM (default) | SM (relax) | SM (default) | SM (relax) | OMGS |  |
| contig/scaffold N50 (bp) | 5,633,882 | 14,354,752 | 14,354,752 | 13,527,997 | 14,892,235 | **16,364,046** |  |
| contig/scaffold L50 | 28 | 14 | 14 | 14 | 13 | 12 |  |
| total assembled (bp) | 511,101,122 | 523,520,329 | 523,139,705 | 521,540,185 | 525,105,345 | 520,697,623 |  |
| # contigs/scaffolds | 948 | 823 | 824 | 817 | 814 | 805 |  |
| # contigs/scaffolds $\geq 100$kb | 269 | 150 | 151 | 146 | 143 | 139 | |
| # contigs/scaffolds $\geq 1$Mbp | 94 | 48 | 48 | 48 | 46 | 46 | |
| # contigs/scaffolds $\geq 10$Mbp | 10 | 21 | 21 | 21 | 22 | 21 | |
| contigs/scaffolds with consistent LG (bp) | 425,812,490 | 402,344,751 | 401,964,127 | 420,269,616 | 403,637,207 | **431,921,182** |  |

As said, the three scaffolding tools were run on a chimera-free assembly of cowpea described above using two available Bionano Genomics optical maps (the first obtained using the BspQI nicking enzyme, and the second obtained with the BssSI nicking enzyme). Since SewingMachine can only use a single optical map, we alternated the optical maps in input (BspQI map first, then BssSI and vice versa). SewingMachine provides two outputs depending on the minimum allowed alignment confidence, namely ‘default’ and ‘relax’. Mode ‘relax’ considers more alignments than ‘default’, but it has a higher chance of introducing mis-joins. HybridScaffold failed on the BssSI map, so we could not test it on alternating maps.

Table 1 shows that when using a single optical map, OMGS can generate comparable or better scaffolds than SewingMachine and HybridScaffold. With two optical maps, OMGS’ correctness (“contigs/scaffolds with 100% consistent LG”) and contiguity (N50) are significantly better than other two tools. Observe that OMGS’ correctness (“contigs/scaffolds with 100% consistent LG”) is even better than the input assembly. This can happen when contigs with SNPs belonging to same linkage group are scaffolded with contigs that have no SNP.

We also compared the performance of OMGS, SewingMachine and HybridScaffold when using optical maps corrected by Chimericognizer (on the same cowpea assembly). Observe in Table 2 that OMGS, SewingMachine and HybridScaffold increased the correctness but decreased the contiguity when the corrected BspQI optical map was used. The results on the corrected BssSI optical map or both corrected optical maps did not change significantly. But again, OMGS produced better scaffolds than SewingMachine and HybridScaffold.

### 4.2 Experimental results on *D. melanogaster*

*D. melanogaster* has four pairs of chromosomes: three autosomes, and one pair of sex chromosomes. The fruit fly’s genome is about 139.5 Mb. We downloaded three *D. melanogaster* assemblies generated in [36] (https://github.com/danrdanny/Nanopore_ISO1). The first assembly (295 contigs, total size 141 Mb, N50 = 3 Mb) was generated using Canu [3, 18] on Oxford Nanopore (ONT) reads longer than 1kb. The second assembly (208 contigs, total size 132 Mb, N50 = 3.9 Mb) was generated using MiniMap and MiniAsm [19] using only ONT reads. The third assembly (339 contigs, total size 134 Mb, N50 = 10 Mb) was generated by Platanus [16] and Dbg2Olc [39] using 67.4x of Illumina paired-end reads and the longest 30x ONT reads. The first and third assemblies were polished using NANOPOLISH [21] and Pilon [38]. The Bionano optical for *D. melanogaster* map was provided by the authors of [36]. This BspQI optical map (363 molecules, total size = 246 Mb, N50 = 841 kb) was created using IrysSolve 2.1 from 78,397 raw Bionano molecules (19.9 Gb of data with a mean read length 253 kb).

As said, all tools were run with default parameters, with the exception of OMGS’ minimum confidence, which was set at 20 (default is 15). To evaluate the performance of OMGS, Hybrid-Scaffold and SewingMachine, we compared their output scaffolds to the high-quality reference genome of *D. melanogaster* (release 6.21, downloaded from FlyBase). We reported the total length of correct/non-chimeric scaffolds as a measure of the overall correctness. To determine which scaffolds were incorrect/chimeric we first selected BLAST alignments of the scaffolds against the reference genome which had an e-value lower than 1e-50 and an alignment length higher than 30 kbp. We defined a scaffold *S* to be *chimeric* if *S* had at least two high-quality alignments which satisfied one or more of the following conditions: (i) *S* aligned to different chromosomes; (ii) the orientation of *S*’s alignments were different; or (iii) the difference between the distance of alignments on the scaffold and the distance of alignments on the reference sequence was larger than 100 Kbp.

Table 3 reports the main statistics for the three *D. melanogaster* scaffolded assemblies. Even with one map, OMGS’ scaffolds are better than SewingMachine and HybridScaffold.

**Table 3:** Comparing OMGS, SewingMachine (SM) and HybridScaffold (HS) on three *D. melanogaster* assemblies (produced by MiniA,sm, Canu, and Dbg2Olc) using the BspQI optical map. Numbers in boldface highlight the best N50 and the best scaffold consistency with the reference genome

|  | MINIASM assembly |  |  |  |  |
| --- | --- | --- | --- | --- | --- |
|  | Input | SM (default) | SM (relax) | HS | OMGS |
| contig/scaffold N50 (bp) | 3,866,686 | 4,494,241 | **4,906,224** | 3,866,686 | **4,906,224** |
| contig/scaffold L50 | 9 | 8 | 8 | 9 | 8 |
| total assembled (bp) | 131,856,353 | 132,480,826 | 133,233,999 | 132,138,056 | 132,838,677 |
| # contigs/scaffolds | 208 | 205 | 203 | 206 | 206 |
| # contigs/scaffolds $\geq 100$kbp | 85 | 82 | 80 | 83 | 83 |
| # contigs/scaffolds $\geq 1$Mbp | 26 | 26 | 25 | 26 | 25 |
| # contigs/scaffolds $\geq 10$Mbp | 2 | 2 | 2 | 2 | 2 |
| non-chimeric contigs/scaffolds (bp) | 131,317,873 | 125,305,638 | **132,695,519** | 131,174,201 | 132,300,197 |

|  | CANU assembly |  |  |  |  |
| --- | --- | --- | --- | --- | --- |
|  | Input | SM (default) | SM (relax) | HS | OMGS |
| contig/scaffold N50 (bp) | 3,004,953 | 3,004,953 | 3,004,953 | 3,918,649 | **5,336,340** |
| contig/scaffold L50 | 11 | 11 | 11 | 10 | 7 |
| total assembled (bp) | 140,720,404 | 140,923,974 | 140,923,974 | 140,867,226 | 140,960,395 |
| # contigs/scaffolds | 295 | 291 | 291 | 286 | 280 |
| # contigs/scaffolds $\geq 100$kbp | 111 | 107 | 107 | 102 | 96 |
| # contigs/scaffolds $\geq 1$Mbp | 31 | 31 | 31 | 29 | 27 |
| # contigs/scaffolds $\geq 10$Mbp | 1 | 1 | 1 | 1 | 5 |
| non-chimeric contigs/scaffolds (bp) | 140,720,404 | 140,923,974 | 140,923,974 | 140,867,226 | **140,960,395** |

|  | DBG2OLC assembly |  |  |  |  |
| --- | --- | --- | --- | --- | --- |
|  | Input | SM (default) | SM (relax) | HS | OMGS |
| contig/scaffold N50 (bp) | 10,113,899 | 11,223,142 | 11,223,142 | 12,785,467 | **12,928,771** |
| contig/scaffold L50 | 6 | 5 | 5 | 5 | 4 |
| total assembled (bp) | 134,109,164 | 134,164,629 | 134,164,629 | 134,162,857 | 134,208,377 |
| # contigs/scaffolds | 339 | 337 | 337 | 331 | 327 |
| # contigs/scaffolds $\geq 100$kbp | 78 | 76 | 76 | 70 | 66 |
| # contigs/scaffolds $\geq 1$Mbp | 22 | 22 | 22 | 17 | 16 |
| # contigs/scaffolds $\geq 10$Mbp | 6 | 6 | 6 | 5 | 7 |
| non-chimeric contigs/scaffolds (bp) | 134,109,164 | 134,164,629 | 134,164,629 | 134,162,857 | **134,208,377** |

## 5. Conclusions

We presented a scaffolding tool called OMGS for improving the contiguity of *de novo* genome assembly using one or multiple optical maps. OMGS solves several optimization problems to generate scaffolds with optimal contiguity and correctness. Experimental results on *V. unguiculata* and *D. melanogaster* clearly demonstrate that OMGS outperforms SewingMachine and HybridScaffold both in contiguity and correctness using multiple optical maps.

## Acknowledgements

This work was supported in part by National Science Foundation grants IIS-1814359, IOS-1543963, IIS-1526742 and IIS-1646333, the Natural Science Foundation of China grant 61772197 and the National Key Research and Development Program of China grant 2018YFC0910404.

## Appendix

### A. DAG unique ordering

#### Algorithm 1

~~~
Sketch of the algorithm for checking whether a DAG provides an unique ordering
1: **procedure** Order Uniqueness Check(*G* = (*V, E*))
2:     *S* = nodes with no incoming edges
3:     **while** *S ≠* ∅ **do**
4:        **if** |*S*| *>* 1 **then**
5:          **return** *False*
6:        remove a node *n* from *S*
7:        **for each** node *m* with an edge *e* = (*n, m*) **do**
8:             remove edge *e* from the *E*
9:             **if** *m* has no other incoming edges **then**
10:                  insert *m* into *S*
11: **return** *True*
~~~

### B. Statistical test for repetitive regions

Here we provide additional details for the estimation of *σ*^2^ during the analysis of repetitive regions. Recall that we collect all estimated repetitive lists in set *R* = {*D*_*p*_ is estimated repetitive|*p* = 1, *…, P*} and the estimated mean 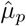 for each distance list *D*_*p*_ in the set *R*, where *P* is the total number of estimated repetitive lists. For each *D*_*p*_, the distances *d*_*i*_’s are distributed as a Gaussian with mean 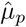 and variance *σ*^2^. According to the density function of Gaussian distribution, the log likelihood of one *D*_*p*_ is

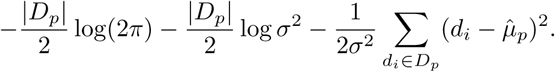

The total log likelihood is the sum of the log likelihoods across all *D*_*p*_’s in *R*, which is

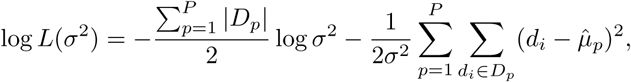

after ignoring all terms not related to *σ*^2^. To maximize log *L*(*σ*^2^), we require that the derivative of total log likelihood

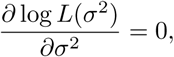

that is,

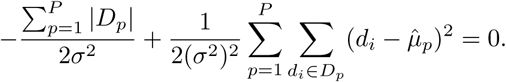

After some simplification, the estimator for variance becomes

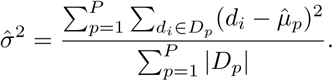

### C. Density function of *d*_*max*_ − *d*_*min*_

Here we provide additional details for calculating the density function of *d*_*max*_ − *d*_*min*_. It is well-known that the joint density function of order statistics is

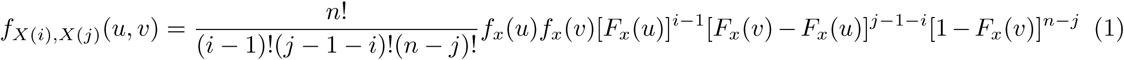

for −*∞ < u < v <* +*∞*, where *X*(*i*) and *X*(*j*) are the *i*-th and *j*-th order statistics in *X*_1_, *…, X*_*n*_ and *F*_*x*_ and *f*_*x*_ are the distribution function and density function of each *X*_*i*_, respectively. Using (1), the joint density function of (*d*_*max*_,*d*_*min*_) can be expressed as

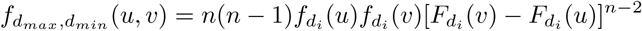

for −*∞< u < v <* +*∞*, where 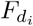 and 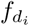 are the distribution function and density function of 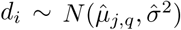, respectively.

Now, let *X* = *d*_*max*_ − *d*_*min*_ and *Y* = *d*_*min*_. Then *d*_*max*_ = *X* + *Y* and *d*_*min*_ = *Y*, and the corresponding Jacobian determinant is

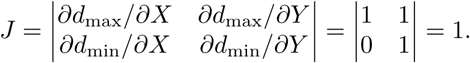

Thus, the joint density function of (*X, Y*) is given by

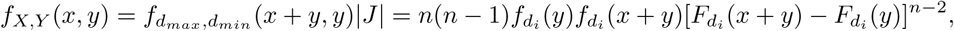

where *x ≥* 0 and −*∞ < y <* +*∞*. By integrating over *Y*, the density function of *X* = *d*_*max*_ − *d*_*min*_ becomes

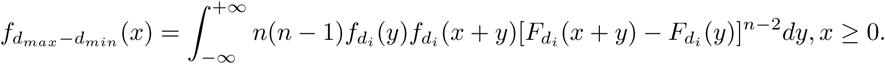

### D. Gap estimation

Here we provide additional details for calculating the log likelihood function when estimating gaps. Recall that *l*_*p*_, *…, l*_*q-*1_ are independent chi-square random variables, and 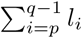 is chi-square distributed with degree of freedom 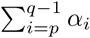. Since the density function of a chi-square random variable *X* with degree of freedom *k* is

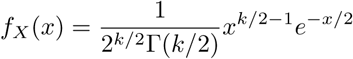

where Γ is the gamma function, the likelihood of 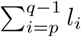 with observation

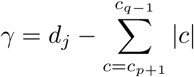

is

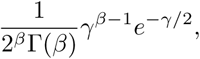

where 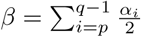. Therefore, the log likelihood function for one sample is

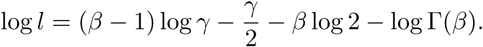

The total log likelihood is the sum of the log likelihoods across all samples.

## Footnotes

1 length for which the set of contigs/scaffolds of that length or longer accounts for at least half of the assembly size

2 minimum number of contigs/scaffolds accounting for at least half of the assembly

## References

[1] Mordecai Avriel. Nonlinear programming: analysis and methods. Courier Corporation, 2003.

[2] Ali Baharev, Hermann Schichl, Arnold Neumaier, and TOBIAS Achterberg. An exact method for the minimum feedback arc set problem. University of Vienna, 10:35–60, 2015.

[3] Konstantin Berlin, Sergey Koren, Chen-Shan Chin, James P Drake, Jane M Landolin, and Adam M Phillippy. Assembling large genomes with single-molecule sequencing and locality-sensitive hashing. Nature biotechnology, 33(6):623, 2015.

[4] Derek M Bickhart, Benjamin D Rosen, Sergey Koren, Brian L Sayre, Alex R Hastie, Saki Chan, Joyce Lee, Ernest T Lam, Ivan Liachko, Shawn T Sullivan, et al. Single-molecule sequencing and chromatin conformation capture enable de novo reference assembly of the domestic goat genome. Nature genetics, 49(4):643, 2017.

[5] Marten Boetzer, Christiaan V Henkel, Hans J Jansen, Derek Butler, and Walter Pirovano. Scaffolding pre-assembled contigs using sspace. Bioinformatics, 27(4):578–579, 2010.

[6] Chen-Shan Chin, Paul Peluso, Fritz J Sedlazeck, Maria Nattestad, Gregory T Concepcion, Alicia Clum, Christopher Dunn, Ronan O’Malley, Rosa Figueroa-Balderas, Abraham Morales-Cruz, et al. Phased diploid genome assembly with single-molecule real-time sequencing. Nature methods, 13(12):1050, 2016.

[7] Nicolas Daccord, Jean-Marc Celton, Gareth Linsmith, Claude Becker, Nathalie Choisne, Elio Schijlen, Henri van de Geest, Luca Bianco, Diego Micheletti, Riccardo Velasco, et al. High-quality de novo assembly of the apple genome and methylome dynamics of early fruit development. Nature genetics, 49(7):1099, 2017.

[8] Adel Dayarian, Todd P Michael, and Anirvan M Sengupta. Sopra: Scaffolding algorithm for paired reads via statistical optimization. BMC bioinformatics, 11(1):345, 2010.

[9] Erik D Demaine and Nicole Immorlica. Correlation clustering with partial information. In Approximation, Randomization, and Combinatorial Optimization.. Algorithms and Techniques, pages 1–13. Springer, 2003.

[10] Anders Dessmark, Jesper Jansson, Andrzej Lingas, Eva-Marta Lundell, and Mia Persson. On the approximability of maximum and minimum edge clique partition problems. International Journal of Foundations of Computer Science, 18(02):217–226, 2007.

[11] Nilgun Donmez and Michael Brudno. Scarpa: scaffolding reads with practical algorithms. Bioinformatics, 29(4):428–434, 2012.

[12] Song Gao, Niranjan Nagarajan, and Wing-Kin Sung. Opera: reconstructing optimal genomic scaffolds with high-throughput paired-end sequences. In International Conference on Research in Computational Molecular Biology, pages 437–451. Springer, 2011.

[13] Alexey A Gritsenko, Jurgen F Nijkamp, Marcel JT Reinders, and Dick de Ridder. Grass: a generic algorithm for scaffolding next-generation sequencing assemblies. Bioinformatics, 28(11):1429–1437, 2012.

[14] Martin Hunt, Chris Newbold, Matthew Berriman, and Thomas D. Otto. A comprehensive evaluation of assembly scaffolding tools. Genome Biology, 15(3):R42, Mar 2014.

[15] Wen-Biao Jiao, Gonzalo Garcia Accinelli, Benjamin Hartwig, Christiane Kiefer, David Baker, Edouard Severing, Eva-Maria Willing, Mathieu Piednoel, Stefan Woetzel, Eva Madrid-Herrero, Bruno Huettel, Ulrike Hümann, Richard Reinhard, Marcus A Koch, Daniel Swan, Bernardo Clavijo, George Coupland, and Korbinian Schneeberger. Improving and correcting the contiguity of long-read genome assemblies of three plant species using optical mapping and chromosome conformation capture data. Genome Res., 27(5):778–786, May 2017.

[16] Rei Kajitani, Kouta Toshimoto, Hideki Noguchi, Atsushi Toyoda, Yoshitoshi Ogura, Miki Okuno, Mitsuru Yabana, Masahira Harada, Eiji Nagayasu, Haruhiko Maruyama, et al. Efficient de novo assembly of highly heterozygous genomes from whole-genome shotgun short reads. Genome research, pages gr–170720, 2014.

[17] Sergey Koren, Todd J Treangen, and Mihai Pop. Bambus 2: scaffolding metagenomes. Bioinformatics, 27(21):2964–2971, 2011.

[18] Sergey Koren, Brian P Walenz, Konstantin Berlin, Jason R Miller, Nicholas H Bergman, and Adam M Phillippy. Canu: scalable and accurate long-read assembly via adaptive k-mer weighting and repeat separation. Genome research, pages gr–215087, 2017.

[19] Heng Li. Minimap and miniasm: fast mapping and de novo assembly for noisy long sequences. Bioinformatics, 32(14):2103–2110, 2016.

[20] Yu Lin, Jeffrey Yuan, Mikhail Kolmogorov, Max W Shen, Mark Chaisson, and Pavel A Pevzner. Assembly of long error-prone reads using de bruijn graphs. Proceedings of the National Academy of Sciences, 113(52):E8396–E8405, 2016.

[21] Nicholas J Loman, Joshua Quick, and Jared T Simpson. A complete bacterial genome assembled de novo using only nanopore sequencing data. Nature methods, 12(8):733, 2015.

[22] Ruibang Luo, Binghang Liu, Yinlong Xie, Zhenyu Li, Weihua Huang, Jianying Yuan, Guangzhu He, Yanxiang Chen, Qi Pan, Yunjie Liu, Jingbo Tang, Gengxiong Wu, Hao Zhang, Yujian Shi, Yong Liu, Chang Yu, Bo Wang, Yao Lu, Changlei Han, David W Cheung, Siu-Ming Yiu, Shaoliang Peng, Zhu Xiaoqian, Guangming Liu, Xiangke Liao, Yingrui Li, Huanming Yang, Jian Wang, Tak-Wah Lam, and Jun Wang. Soapdenovo2: an empirically improved memory-efficient short-read de novo assembler. GigaScience, 1(1):18–18, 12 2012.

[23] Martin Mascher, Heidrun Gundlach, Axel Himmelbach, Sebastian Beier, Sven O Twardziok, Thomas Wicker, Volodymyr Radchuk, Christoph Dockter, Pete E Hedley, Joanne Russell, et al. A chromosome conformation capture ordered sequence of the barley genome. Nature, 544(7651):427, 2017.

[24] María Muñoz-Amatriaín, Hamid Mirebrahim, Pei Xu, Steve I Wanamaker, MingCheng Luo, Hind Alhakami, Matthew Alpert, Ibrahim Atokple, Benoit J Batieno, Ousmane Boukar, et al. Genome resources for climate-resilient cowpea, an essential crop for food security. The Plant Journal, 89(5):1042–1054, 2017.

[25] Niranjan Nagarajan, Timothy D Read, and Mihai Pop. Scaffolding and validation of bacterial genome assemblies using optical restriction maps. Bioinformatics, 24(10):1229–1235, 2008.

[26] Weihua Pan and Stefano Lonardi. Accurate detection of chimeric contigs via bionano optical maps. Bioinformatics, 2018.

[27] Weihua Pan, Steve I Wanamaker, Audrey MV Ah-Fong, Howard S Judelson, and Stefano Lonardi. Novo&stitch: accurate reconciliation of genome assemblies via optical maps. Bioinformatics, 34(13):143–151, 2018.

[28] Matthew Pendleton, Robert Sebra, Andy Wing Chun Pang, Ajay Ummat, Oscar Franzen, Tobias Rausch, Adrian M Stütz, William Stedman, Thomas Anantharaman, Alex Hastie, et al. Assembly and diploid architecture of an individual human genome via single-molecule technologies. Nature methods, 12(8):780, 2015.

[29] Mihai Pop, Daniel S Kosack, and Steven L Salzberg. Hierarchical scaffolding with bambus. Genome research, 14(1):149–159, 2004.

[30] Subrata Saha and Sanguthevar Rajasekaran. Efficient and scalable scaffolding using optical restriction maps. BMC genomics, 15(5):S5, 2014.

[31] Leena Salmela, Veli Mäkinen, Niko Välimäki, Johannes Ylinen, and Esko Ukkonen. Fast scaffolding with small independent mixed integer programs. Bioinformatics, 27(23):3259–3265, 2011.

[32] Akhtar Samad, EF Huff, Weiwen Cai, and David C Schwartz. Optical mapping: a novel, single-molecule approach to genomic analysis. Genome research, 5(1):1–4, 1995.

[33] Jennifer M Shelton, Michelle C Coleman, Nic Herndon, Nanyan Lu, Ernest T Lam, Thomas Anantharaman, Palak Sheth, and Susan J Brown. Tools and pipelines for bionano data: molecule assembly pipeline and fasta super scaffolding tool. BMC genomics, 16(1):734, 2015.

[34] Jared T Simpson and Richard Durbin. Efficient de novo assembly of large genomes using compressed data structures. Genome research, 22(3):549–556, 2012.

[35] Jared T Simpson, Kim Wong, Shaun D Jackman, Jacqueline E Schein, Steven JM Jones, and Inanç Birol. Abyss: a parallel assembler for short read sequence data. Genome research, pages gr–089532, 2009.

[36] Edwin A Solares, Mahul Chakraborty, Danny E Miller, Shannon Kalsow, Kate E Hall, Anoja G Perera, JJ Emerson, and R Scott Hawley. Rapid low-cost assembly of the drosophila melanogaster reference genome using low-coverage, long-read sequencing. bioRxiv, page 267401, 2018.

[37] Haibao Tang, Xingtan Zhang, Chenyong Miao, Jisen Zhang, Ray Ming, James C Schnable, Patrick S Schnable, Eric Lyons, and Jianguo Lu. Allmaps: robust scaffold ordering based on multiple maps. Genome biology, 16(1):3, 2015.

[38] Bruce J Walker, Thomas Abeel, Terrance Shea, Margaret Priest, Amr Abouelliel, Sharadha Sakthikumar, Christina A Cuomo, Qiandong Zeng, Jennifer Wortman, Sarah K Young, et al. Pilon: an integrated tool for comprehensive microbial variant detection and genome assembly improvement. PloS one, 9(11):e112963, 2014.

[39] Chengxi Ye, Christopher M Hill, Shigang Wu, Jue Ruan, and Zhanshan Sam Ma. Dbg2olc: efficient assembly of large genomes using long erroneous reads of the third generation sequencing technologies. Scientific reports, 6:31900, 2016.

[40] Jie Zheng and S. Lonardi. Discovery of repetitive patterns in dna with accurate boundaries. In Fifth IEEE Symposium on Bioinformatics and Bioengineering (BIBE’05), pages 105–112, Oct 2005.

